# *Drosophila* re-zero their path integrator at the center of a fictive food patch

**DOI:** 10.1101/2021.01.18.427191

**Authors:** Amir H. Behbahani, Emily H. Palmer, Román A. Corfas, Michael H. Dickinson

**Affiliations:** Division of Biology & Bioengineering, California Institute of Technology, Pasadena CA 91125, USA

**Keywords:** *Drosophila*, path integration, odometry, place memory, state-dependent models

## Abstract

The ability to keep track of one’s location in space is a critical behavior for animals navigating to and from a salient location, and its computational basis is now beginning to be unraveled. Here, we tracked flies in a ring-shaped channel as they executed bouts of search triggered by optogenetic activation of sugar receptors. Unlike experiments in open field arenas, which produce highly tortuous search trajectories, our geometrically constrained paradigm enabled us to monitor flies’ decisions to move toward or away from the fictive food. Our results suggest that flies use path integration to remember the location of a food site even after it has disappeared, and that flies can remember the location of a former food site even after walking around the arena one or more times. To determine the behavioral algorithms underlying *Drosophila* search, we developed multiple state transition models and found that flies likely accomplish path integration by combining odometry and compass navigation to keep track of their position relative to the fictive food. Our results indicate that whereas flies re-zero their path integrator at food when only one feeding site is present, they adjust their path integrator to a central location between sites when experiencing food at two or more locations. Together, this work provides a simple experimental paradigm and theoretical framework to advance investigations of the neural basis of path integration.

## INTRODUCTION

For many animals, including humans, the ability to return to a specific location such as a nest or food resource is essential for survival^1^. One strategy for revisiting a specific location is to use external cues such as chemical signals or visual landmarks^2–4^. Another strategy that works in visually poor landscapes or featureless environments^5, 6^ is to perform path integration, that is, to cumulatively integrate along a path, using a measure of distance traveled (odometry) and body orientation in the direction of travel (heading), thus making it possible to calculate a direct path between any current position and a starting point. Since Darwin first suggested that animals might perform path integration to navigate between food and their nests^7^, ample evidence has emerged that many animals employ this strategy. The behavior has been best characterized in ants and bees^8–10^ but has been identified in many species including mantis shrimps^11^, bats^12^, dogs^13^, and rats^14^. Whereas the entire process of path integration is difficult to observe directly, it is often manifest in the act of homing, when an animal executes a straight path (a so-called ‘home run’) back to its nest after completing a tortuous excursion in search of food^15–18^. Animals can also walk directly from their nest to a food site after their first visit to that location^19–21^. In all these classic cases with virtuosic path integrators such as ants and honeybees, animals reset their path integrator to zero at their nest or hive location. However, animals without a nest or hive may instead zero their path integrator at a food site, or at the center of a cluster of food sites.

Path integration can operate on the scale of hundreds of meters, as exemplified by desert ants^15^, or many kilometers, as in bees^8^; however, it can also occur over much smaller spatial scales. In ants, for example, homing is often accompanied by a local search when the forager arrives near the nest, but not near enough to immediately find it^18, 22–24^. Although seemingly random, these local searches are structured and centered, suggesting the animal is keeping track of its best estimate of the nest’s location. Such local searches are not restricted to central place foragers such as ants and bees; for example, hungry blowflies execute local searches in the vicinity of small food items they have sampled, a behavior that Vincent Dethier described as a ‘dance’^25^. *Drosophila melanogaster* also exhibit these local searches near small spots of food^26, 27^. Optogenetic activation of sugar receptors can substitute for the presence of actual food in initiating this behavior^27, 28^. Fruit flies can perform this food-centered search in the absence of external stimuli or landmarks, indicating that they can rely on idiothetic (internal) information to keep track of their location^27^. These local searches consist of highly tortuous trajectories in which it is difficult to classify instances when the fly is walking either directly away or toward the food site, e.g., executing a so-called ‘home run’, making analysis of the behavior quite challenging.

In this study, we deliberately constrained the two-dimensional motion of flies by confining them to an annular channel. In this constrained arena, local searches consist of back-and-forth runs centered around arbitrarily defined food zones where the flies receive or have recently received optogenetic activation of sugar receptors. Due to its geometric simplicity, our arena allowed us to address several key questions that are difficult to test in an open field arena. By analyzing the flies’ behavior after they walked in one or more complete circles around the arena, we provide strong evidence that *Drosophila melanogaster* are capable of two-dimensional, idiothetic path integration to search in the vicinity of a previously exploited food site. In addition, we were able to examine how an animal centers its search excursions when offered a cluster of locations. Our results suggest that rather than using the location of the most recently visited food location to zero their path integrator, flies are able to center their search at a single location within a patch of food.

## RESULTS

### Flies remember the position of a single food site

To investigate the behavioral algorithms underlying path integration, we tracked individual flies as they performed local search in an annular arena in which the flies were constrained to walking within a circular channel (Figure 1A). Using an automated closed-loop system, we optogenetically activated sugar-sensing neurons whenever the flies (*Gr5a-GAL4>UAS-CsChrimson*) occupied a designated, featureless, food zone. The one-second optogenetic light pulse triggered by residence in the food zone was followed by a 15-second refractory period during which time the stimulus was kept off, regardless of the fly’s position. Aside from these brief optogenetic pulses, all experiments were conducted in complete darkness. For convenience, we sometimes refer to optogenetic food zones as ‘food’, and optogenetic activation events as ‘food stimuli’, although in no cases did the animals experience actual food.

**Figure 1.**
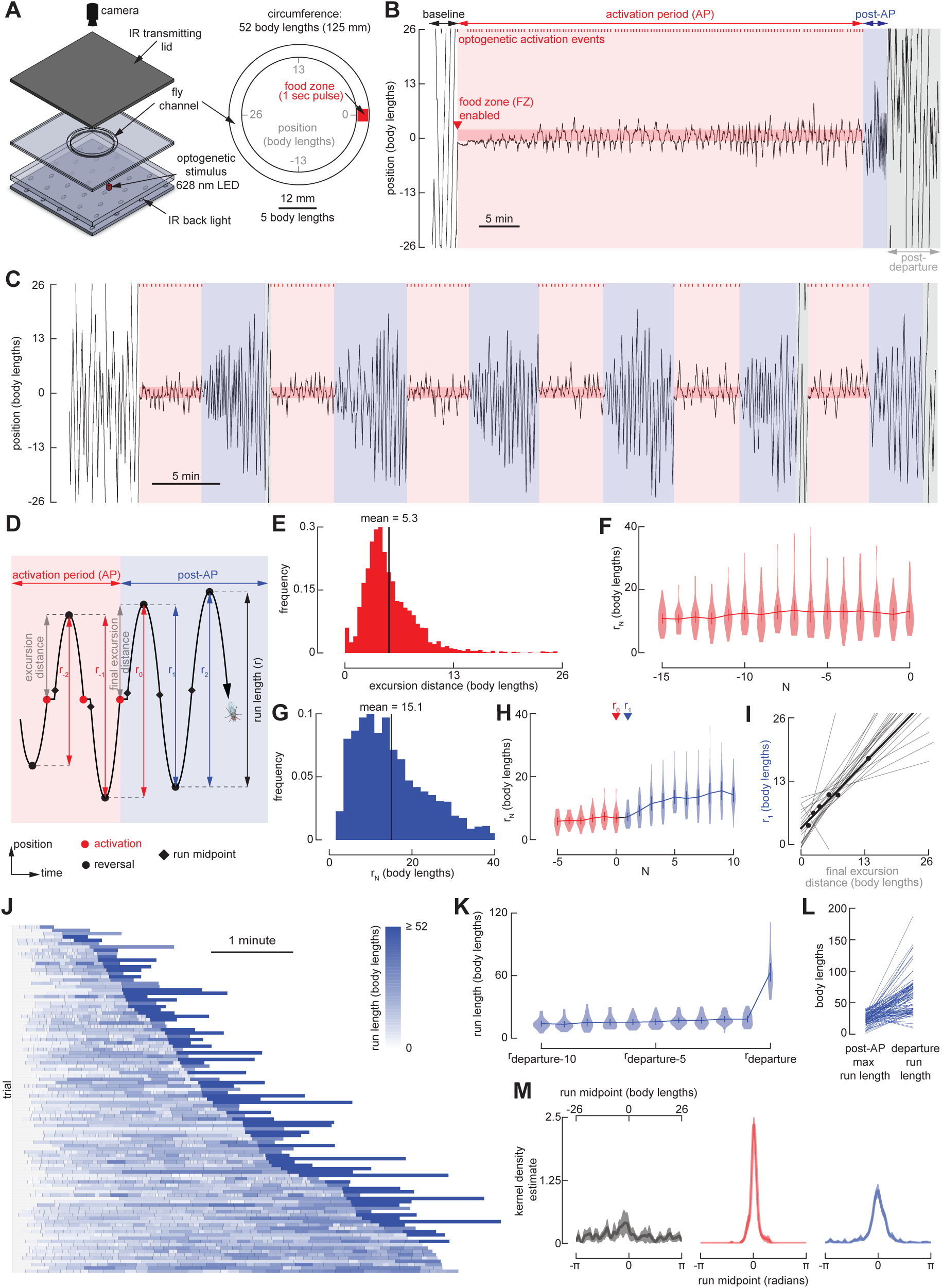
Repeated back-and-forth excursions constitute a local search around a fictive food location. **(A)** Schematic of the experimental setup (left) and annular arena (right). An overhead camera tracks the position of an individual *Gr5a-GAL4>UAS-CsChrimson* female fly, in real-time, as it explores a 4 mm-wide circular channel, ∼52 body lengths (BL) in circumference. Whenever the fly occupies the featureless food zone, it receives a one-second pulse of optogenetic activation of sugar-sensing neurons via a 628 nm LED positioned beneath the channel, followed by a 15-second refractory period during which the fly cannot receive activation. An infrared (IR) backlight and IR-transmitting lid enable behavioral tracking while otherwise maintaining complete darkness for the fly aside from the brief optogenetic pulses. **(B)** Example fly trajectory. To simplify the display and analysis of the data, we transformed the curved trajectories of the flies in the circular channel into a wrapped one-dimensional path. This experiment begins with a baseline period, during which the fly does not receive optogenetic activation, followed by a 40-minute activation period (AP, red) during which the optogenetic protocol is operational, followed by a post-activation-period (post-AP, blue) during which the optogenetic protocol is switched off. The post-AP is defined as ending when the fly executes its first run straying more than 26 body lengths (i.e., ½ the arena perimeter) from the food zone, hereafter termed the ‘departure run’. The remaining trajectory is referred to as post-departure (grey). Optogenetic stimulation events during the AP are indicated as red tick-marks (top). See also Figure S1A. **(C)** As in (B), for an experiment with six serial trials each consisting of a 5-minute AP followed by a 5-minute post-AP. See also Figure S1B and Video S1. **(D)** Schematic, showing features of local search. After encountering a food stimulus during the AP, flies walk a given excursion distance (grey), reverse direction, and perform a run back towards the food. The distance between two consecutive reversals is a run length (*r*), where *r_0_* is the run length between the final reversal of the AP and the first reversal of the post-AP, and all other runs are numbered with respect to *r_0_*. The run midpoint is defined as the halfway point between two consecutive reversals. **(E)** Distribution of all excursion distances, from the 40-min AP experiments as in (B). (N = 29 flies, 2494 food excursions). **(F)** Run lengths for the final 16 runs of the AP, including *r_0_*, from the 40-min AP experiments. Data from trials with fewer than 16 AP runs are included. (N = 29 flies). Throughout the paper, error bars depict 95% confidence intervals and violin plots indicate full data distributions. **(G)** Mean distribution of the run lengths of the post-AP from the trial-based experiments as in (C). (N = 22 flies, n = 110 trials). **(H)** Run lengths for the final 6 runs of the AP, including *r_0_*, and the first 10 runs of the post-AP, from the trial-based experiments as in (C). (N = 22 flies, n = 110 trials). Labels indicate the final run of the AP (*r_0_*) and the first run of the post-AP (*r_1_*). **(I)** Relationship between the final excursion distance during the AP and the first run of the post-AP (*r_1_*). Black dots indicate *r_1_* vs. last excursion for 6 trials from the fly in (C), and the black line indicates the linear regression for this fly. Grey lines indicate linear regressions for all remaining flies with data from at least 3 trials. (N = 20 flies). **(J)** Sequences of post-AP runs and their associated departure run, sorted by the duration of the post-AP. Each row corresponds to a single trial from experiments as in (C), where the length of each box corresponds to the duration of each run, and the color of each box indicates run length. (N = 22 flies, n = 110 trials). Note that in 11 trials at the bottom of the panel, the flies did not execute a departure run before the next AP began. **(K)** Run lengths for the final 10 runs of the post-AP, as well as the departure run, from data in (J). Data from trials with fewer than 10 post-AP runs are included. **(L)** Length comparison of the longest post-AP run, and corresponding departure run, for each trial, from data in (J). The 11 trials without a departure run were not included in this analysis. **(M)** Normalized kernel density estimate (KDE) of the run midpoint in baseline (left), AP (middle), and post-AP (right). (N = 22 flies). Throughout the paper, shaded regions indicate 95% confidence interval. Throughout the paper, the KDE is calculated for each fly for ϰ = 200 and then the mean and 95% confidence interval is calculated for the individual fly’s KDE. For post-AP comparison with simple models, with run lengths randomly drawn from either the empirically derived data shown in Figure 1J (excluding the departure runs), or a Lévy distribution fit to the same data, see Figure S2.

Examples of local searches are plotted in Figures 1B and 1C (for additional examples, see Figure S1). To simplify the display and analysis of the data, we have transformed the curved trajectories of the flies in the circular channel into one-dimensional paths. All experiments began with a baseline period, during which the optogenetic protocol was not operational. This baseline period was followed by an activation period (AP), during which the optogenetic protocol was in effect, i.e., the fly received the one-second food stimulus followed by the 15-second refractory period whenever it occupied the food zone. Each AP was followed by a post-activation period (post-AP) during which the optogenetic protocol was suspended such that flies did not receive food stimuli. Some experiments consisted of a 40-minute AP, followed by a single 10-minute post-AP (Figure 1B). Other experiments used a repeating trial structure, in which each trial consisted of a 5-minute AP followed by a 5-minute post-AP (Figure 1C).

In the annular arena, flies can either walk clockwise, walk counterclockwise, pause, or reverse direction. We defined the distance between two consecutive reversals as a ‘run length’, *r* (Figure 1D). During the baseline period, flies continuously explored the entire arena, generally performing long runs interspersed with occasional reversals. During the AP, food stimuli consistently triggered local search excursions typically characterized by a stereotyped sequence of behaviors: upon activation of sugar-receptors, the flies briefly paused, continued to walk a few body lengths away from the food, performed a reversal, returned to the food zone, experienced another food stimulus, and then executed a similar excursion in the opposite direction. This process repeated, producing a persistent zig-zagging search pattern during which the flies explored the channel near the food site, while never straying far in either direction. The excursions were reasonably stereotyped, being approximately 5 body lengths in size (Figure 1E), and did not vary substantially during the AP (Figure 1F). We interpret this behavior to be a one-dimensional version of the two-dimensional local searches that were the subject of studies in *Drosophila*^26–28^ in open field arenas, as well as those originally identified by Vincent Dethier in the blowfly, *Phormia*^25^. The relative consistency of these excursion distances in our apparatus was noteworthy, given that there was no external sensory stimulus associated with the termination of each outward run. This suggests that flies’ nervous systems intrinsically produce a motor pattern that generates excursions of a particular spatial scale, in response to the optogenetic activation that we provided.

The most informative data regarding whether the flies retain a spatial memory of the food location came from the post-AP, when the optogenetic protocol was suspended. Despite no longer receiving food stimuli, flies continued to zig-zag back and forth around the disabled food zone (Figures 1B and 1C, Video S1). These post-AP excursions were, however, longer than the AP excursions (mean = 15.1 body lengths, Figure 1G), and tended to increase in length over time toward a plateau by the fifth post-AP run (Figure 1H). The length of the first post-AP run (*r_1_*) tended to correlate strongly with the final excursion distance on a trial by trial basis (Figure 1I). In other words, in our experiments consisting of six successive trials (Figure 1C), the length of the final excursion from the food was strongly correlated with the length of the first run past the former food site. This strong positive correlation was observed in nearly every fly tested (grey lines in Figure 1I).

We also observed that most flies eventually abandoned their post-AP search after some time. To specifically analyze trajectories during which the fly was performing local search, we defined the post-AP as starting at the conclusion of the AP and ending with the execution of what we classified as a ‘departure run.’ The departure run was defined as the first run after the conclusion of the AP during which the fly strayed 26 or more body lengths away from the food zone, thus reaching or passing the opposite side of the arena. The total duration of the post-AP trajectory varied—some flies abandoned the food after 1-2 minutes, while others continued searching for the full 5 minutes of the post-AP (Figure 1J). Regardless of the duration of the post-AP search, the departure run was almost always considerably longer than all the preceding runs (Figures 1K and 1L). In other words, rather than slowly expanding or drifting away from the food site, flies typically terminated the post-AP search by performing an exceptionally long run, perhaps reflecting a change in the fly’s behavioral state.

To derive an estimate of the flies’ spatial memory within the arena, we developed a method by which we measured the midpoint of each run and then convolved each of these locations with a narrow von Mises distribution (κ = 200), the sum of which generated a kernel density estimate (KDE) of where in the arena the fly was focusing its search. The KDE for the baseline (pre-AP) data was flat and noisy, indicating that the flies’ runs were not centered on any particular location within the arena (Figure 1M, left panel). In sharp contrast, the KDEs from the AP and post-AP data were unimodal with a clear peak at the location of the food (Figure 1M, center and right panels). The food-centered peak is expected for the AP, but its existence for the post-AP data is indicative of the flies’ memory of the food site. We will make use of run midpoint KDEs throughout the paper as a means of assessing the flies’ spatial memory under different conditions.

### Flies recognize a former food site after walking completely around the arena

Our analysis of the flies’ search behavior prior to the departure run suggests that they can remember the location of the food, a task they could accomplish using one-dimensional odometry, for example, by simply counting the number of steps taken away from and back toward the food. However, the fact that some flies traveled all the way around the arena after the departure run allowed us to test whether the flies could perform full two-dimensional path integration, a task that would require integrating odometry with their internal compass sense. To test this hypothesis, we conducted experiments in a smaller (∼26 body length circumference) circular channel (Figure 2A) to increase the probability that the flies would walk one or more times around the arena during the post-AP. We exposed flies to six 5-minute APs separated by 5-minute post-APs, in which each pair of APs and post-APs constitutes a single trial. To ensure that flies could not use their own chemical signals or other external features to recognize the food site, we switched the food zone between two locations, spaced 180° apart, for each AP—i.e., during the 1^st^ trial, the food zone was on the right side of the arena, during the 2^nd^ trial, the food zone was on the left side of the arena, etc. (Figure 2A). We classified a post-return period, starting when the fly reached 1 full revolution (26 body length) away from the food site. After seeming to abandon their local search during the post-AP, many flies reinitiated search at the former food site, after completing one or more full revolutions around the arena (Figures 2B-2E, Video S2). The flies’ performance was quite variable in this task, with some flies exhibiting much stronger search behavior after circling the arena than others. Nevertheless, the transit probability averaged across all flies during the post-return period shows clear peaks at the location of the former food zone at integer values of full revolutions within the arena (Figures 2C-E), and the average KDE of the run midpoint distribution exhibited a peak at the former food location (Figure 2F). As described above, these experiments were conducted by changing the position of the food site from one side of the arena to the other in alternate trials, to control for the possibility that flies were simply depositing some chemical cue when they encountered food, which they subsequently used to relocate that position. If this were the case, we would expect that the KDEs would show two peaks, one at each of the alternating food locations. This is not what we found; instead, flies showed no preference for searching at the position of the food zone from the immediately preceding trial (Figure 2G), suggesting that they are not remembering the food site via chemical cues deposited there.

**Figure 2.**
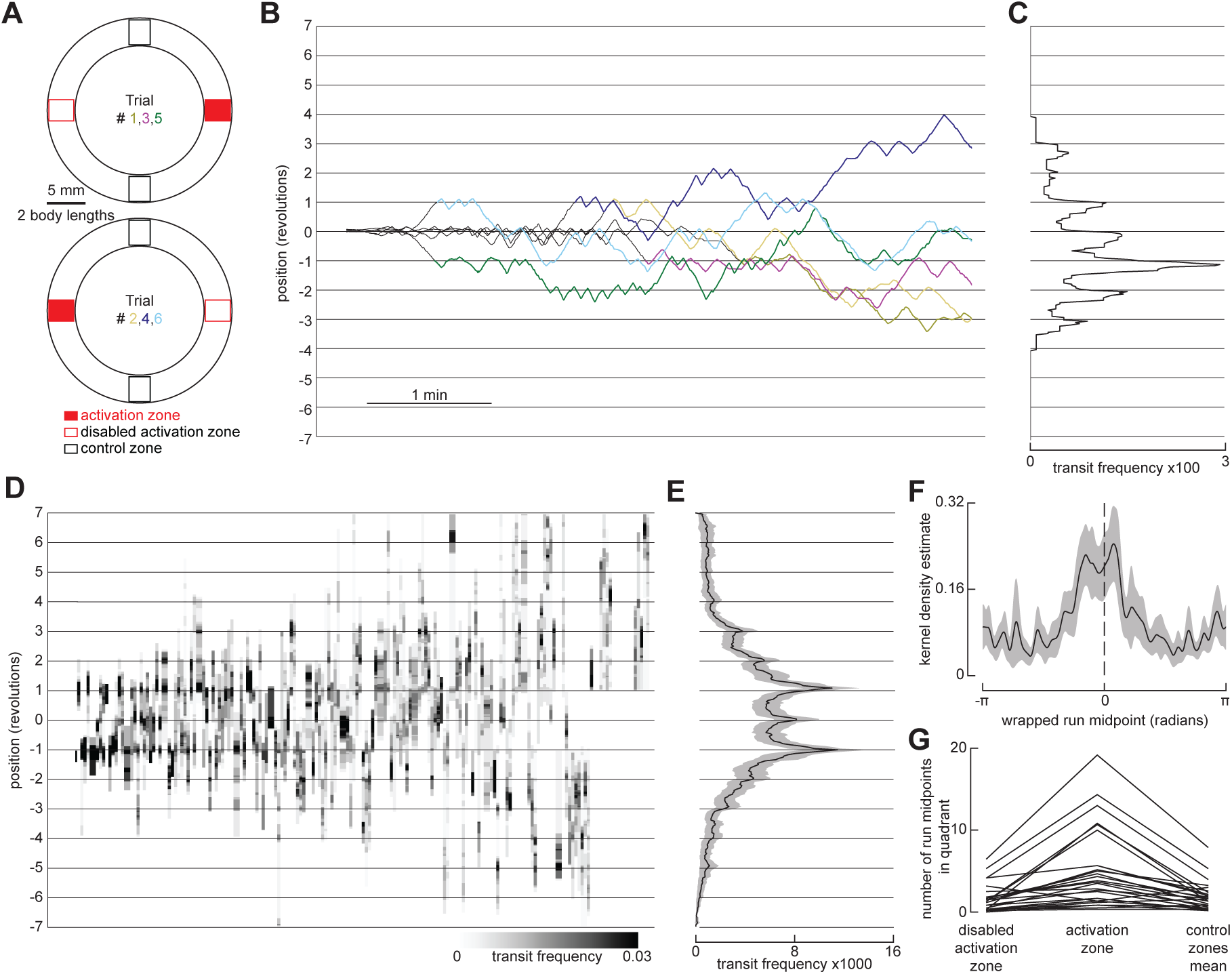
Flies reinitiate a local search at a former fictive food site after circling the arena. **(A)** Schematic of the smaller annular arena (∼26 body lengths), indicating the location of the food zone for each trial, as well as control zones used for analysis. Experiments were done as in Figure 1C, but each food zone was 1.3 body lengths, and the food zone location was alternated from trial to trial. **(B)** Example pre-return (before the fly has circled the arena at least once during the post-AP, grey) and post-return (colored) trajectories from a single experiment where each line corresponds to a single trial and shows the unwrapped trajectory, with gridlines indicating full revolutions around the arena. To align data for analysis, trajectories from even-numbered trials were shifted such that the location of the food zone is always at 0. See also Video S2. For the model recapitulating fly re-initiation of local search at a former fictive food site after circling the arena, see Figure S4 and Video S5. **(C)** Mean distribution of fly transits for post-return trajectories in (B). Transits were calculated using bins 2 BL wide and counted when a fly entered a bin from one side and exited the bin from the other side. **(D)** Heatmap indicating distribution of transits during post-return trajectories, calculated using 4 bins per revolution (dividing the arena into quadrants centered on the food zone, disabled food zone, and each control zone). Each column represents a single trial, with columns sorted by frequency of transits at the 1 or -1 revolution position. (N = 28 flies, n = 168 trials). **(E)** Mean transit distribution for data in (D). **(F)** Normalized kernel density estimate (KDE) of the wrapped run midpoint in the post-return period. (N = 28 flies). **(G)** Number of run midpoints in each arena quadrant during post-return trajectories. (N = 28 flies). Each line shows the values for a single fly, where data from both control quadrants were averaged together.

### An agent-based model recapitulates *Drosophila* local search behavior

To investigate possible algorithms underlying local search, we constructed different agent-based models of the flies’ behavior. The output of each model—a time series of the fly’s position—is generated by a sequence of simulated runs and reversals. First, we tested whether simple models, with run lengths randomly drawn from either the empirically derived data shown in Figure 1J (excluding the departure runs), or a Lévy distribution fit to the same data, could recapitulate the flies’ behavior during the post-AP. In both of these cases, the models failed to produce a sustained, centered, local search; the simulated flies quickly strayed from the trajectory origin (Figure S2). These results suggest that real flies must somehow remember the location of the former food site, and that the centeredness of the post-AP search is not simply a result of starting at the former food location.

To account for the flies’ ability to remember the location of the food, we developed an agent-based, state-transition model (hereafter, food-to-reversal or FR model) that posits the flies’ ability to integrate the distance between the food site and the point at which they reverse direction at the end of each excursion. Figure 3A shows a simple state transition depiction of this model, with a more formal presentation provided in Figure S3B. State-transition diagrams are commonly used in computer science to model systems—self-driving cars, for example^29–31^—where an agent can assume finite states regulated by stochastic or deterministic transitions. In the FR model, flies are initialized in a global search mode and enter a local search mode when they encounter a food stimulus. When in local search mode, simulated flies use odometry to keep track of their distance walked, and when they have completed their target run length, they perform a reversal and select a new target run length as a function of the prior run length. The output of the model—a time series of the fly’s position—captures salient features of the behavior of real flies during both the AP and post-AP periods (Figures 3B-D). To account for the flies’ ability to remember the location of the food site after walking completely around the arena (during the post-return period), the FR model incorporates two orthogonal path integrators and can thus keep track of the food site in two dimensions (Figures 3E-H). This is, in essence, a path integration model, wherein the integrators are set equal to zero upon experience with a food stimulus and a run length is selected. When the fly has deviated a distance from the food site equal to the run length, as measured by the Euclidean norm of the integrators, the fly executes a reversal, zeros its integrators, and selects a new run length as a function of the previous distance walked. The integrators are noiseless, such that the simulated fly has perfect knowledge of its location relative to the food site. We emphasize, however, that this is a purely algorithmic model and we are not asserting whether or not it might be implemented in a neurally plausible manner. The salient feature of the model, however, is that its path integrator is zeroed at the location of the food site and accumulates distance until the fly reverses direction.

**Figure 3.**
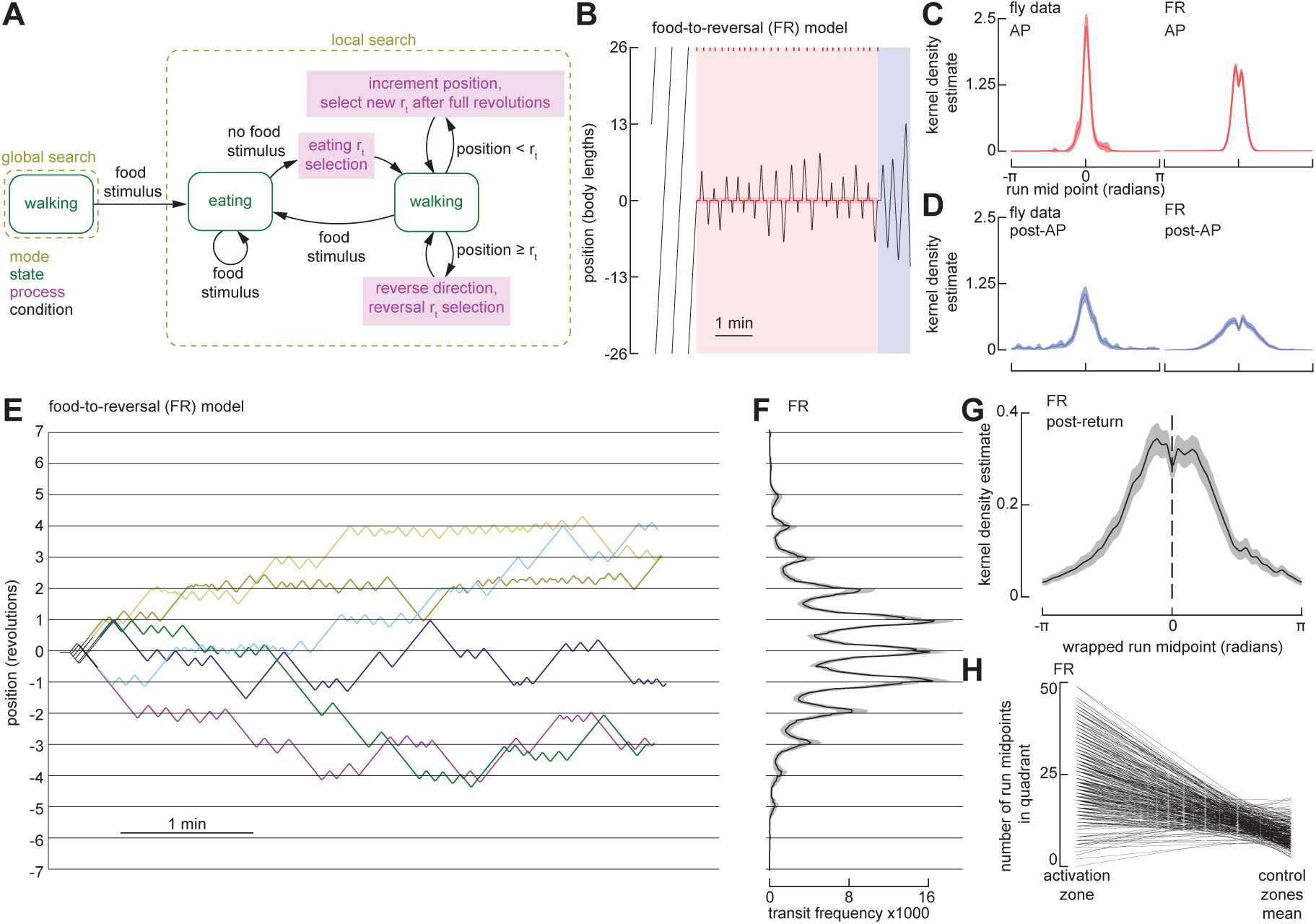
An agent-based model using iterative odometric integration recapitulates *Drosophila* local search around a single fictive food site. **(A)** Schematic of the state-transition diagram for an agent-based model. Arrows indicate transitions—governed by conditions— between search modes, behavioral states, and computational processes. For detailed state transition diagrams, including that of the simulated environment, see Figure S3. **(B)** Example trajectory of FR model simulation in a circular arena with a 52-body length circumference, showing baseline, AP, and post-AP. Plotting conventions as in Figure 1B. **(C)** Normalized kernel density estimate (KDE) of the run midpoint in during AP for flies (left, N = 22 flies) and FR model (right, N = 300). Data for the fly is re-plotted from Figure 1M. **(D)** Normalized kernel density estimate (KDE) of the run midpoint in during post-AP for flies (left, N = 22 flies) and FR model (right, N = 300). Data for the fly is re-plotted from Figure 1M. **(E)** Six representative example trajectories of FR model simulation in a small circular arena with a 26-body length circumference (same as experiments in Figure 2). Plotting conventions as in Figure 2B. **(F)** Mean transit distribution for FR simulations in small arena. (N = 300). Plotting conventions as in Figure 2E. **(G)** Normalized kernel density estimate (KDE) of the wrapped run midpoint in the post-return period for FR simulations in small arena. (N = 300). Plotting conventions as in Figure 2F. **(H)** Number of run midpoints in the food quadrant compared to the other three quadrants during post-return trajectories for FR simulations in small arena. (N = 300). Each line shows the values for a single simulation, where data from all three control quadrants were averaged together.

### Flies expand and recenter their search area to span multiple food sites

We next tested how flies perform local searches within arenas that contain multiple food sites. We modified our annular arena to feature two food zones, separated by 9 body lengths (60°) within the channel. As expected, flies began searching around the first food site they encountered. However, on occasions where the fly encountered the second food site during the course of the search, they often expanded their search area to span both food sites (Figure 4A, Figure S1C, Video S3). We also repeated the two-food experiments with a configuration in which the food was separated by 13 body lengths (90°). Although the results were generally consistent with the data collected with a shorter food separation distance (Figures 4 and S1), it often took the fly longer to find the second food site at the start of the experiment. These experiments with a large separation between food zones underscore the salient phenomenon that the flies’ inward excursions toward the other food zone were substantially longer than their typical outward excursions—an observation that again suggests that their behavior involves some sort of odometric memory that allows them to keep track of the location of two food sites.

**Figure 4.**
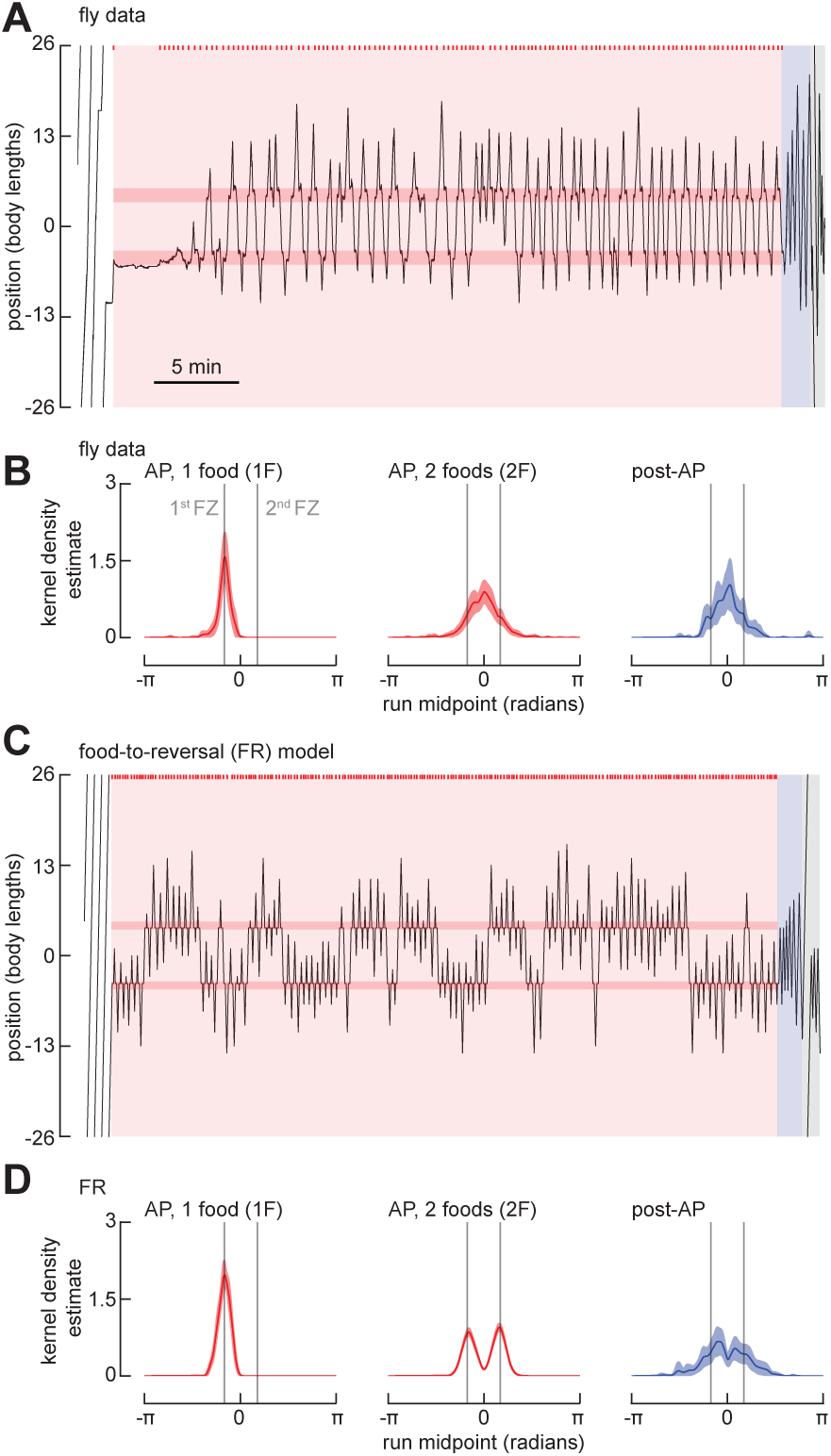
The FR model fails to predict *Drosophila* search behavior around multiple fictive food sites. **(A)** Example trajectory of a fly exploring an annular arena with two food zones, spaced 9 body lengths (BL) apart. The experiment consists of a baseline period, AP, and post-AP. Plotting conventions as in Figure 1B. See also Figure S1C and Video S3. For trajectories of flies exploring an annular arena with two food zones, spaced 13 body lengths (BL) apart, see Figure S1D. **(B)** Normalized KDE of the run midpoint for one-food search (1F, left), the two-food search (trajectory after the fly has encountered the 2^nd^ food zone, 2F, middle), and the post-AP (right). To align data for analysis for 1F, trajectories for flies that found the food located at +4.5 BL first were shifted such that the first food for all flies is -4.5 BL. (N = 29 flies). **(C)** As in (A), for a simulation using the FR model. **(D)** As in (B) for the FR model. The first 300 simulations in which the virtual fly found both food sites are included. (N = 300).

To assess the flies’ behavior during the AP, we segmented the data into the initial segments before they found the second food site, and the remaining segment after they first encountered the second food. We also inverted roughly half of the traces so that all the flies start the AP by foraging around the lower food position (as plotted in Figure 4A). As expected, the run midpoint KDE generated from all the initial segments were centered around the lower food position; however, the KDE generated from the data following discovery of the second food was centered at a location roughly midway between the two food sites, as was the KDE generated from runs during the post-AP (Figure 4B), suggesting that the fly remembers a position midway between the two food sites. We ran our FR model on the same experimental conditions and analyzed the trajectories in the same manner (Figures 4C,D); however, the results failed to replicate the flies’ behavior. During the AP, the simulated flies tended to transition back and forth between local searches around one food site or the other—rarely generating a stable oscillation across the two—and the KDE distribution in the post AP exhibited two peaks (Figure 4D). Thus, while capturing the salient features of flies’ behavior when searching in the vicinity of one food site (Figure 3), the FR model failed to recapitulate the behavior of real flies when searching around two sites (Figure 4).

Based on the failure of the FR model, we developed two distinct models that might explain the flies’ behavior when foraging amid two feeding sites; these are depicted diagrammatically in Figures 5B-C and more formally in Figure S3. In the FR model, experience with any active food zone zeroes all path integrators; the fly therefore integrates from the food site most recently encountered (Figure 5A). At a reversal, the next run length is calculated as the sum of a randomly distributed variable and the integrated value. As the integrated value is only the distance to the nearest food, the search is always centered over one food site rather than the entire food patch (Figure 4C). The first of the two new models, which we call FR’, differs from the simpler FR model most notably in what triggers the zeroing of the path integrators. Whereas the FR model is always zeroed when food is encountered, the FR’ model only zeroes its integrators at the first encounter with a food following a reversal; if a fly experiences a second food site before changing direction, the integrators continue to accumulate (Figure 5B). Thus, when the next run length is calculated at a reversal, the integrated value will take the fly fully back across the entire food patch to the first food site encountered, and the random variable value will ensure the fly continues past that point a short distance. This feature of the FR’ model is able to generate more realistic trajectories in which the simulated fly often zig-zags back and forth across the two food zones (Figure 5D). The FR’ model does not require the fly to zero its path integrators at a location that is not directly associated with a sensory stimulus (e.g., food sensation) or motor action (e.g., run reversal). Further, the FR’ model is intrinsically reflexive and does not require that the fly somehow keep track of the number of food sites visited.

**Figure 5.**
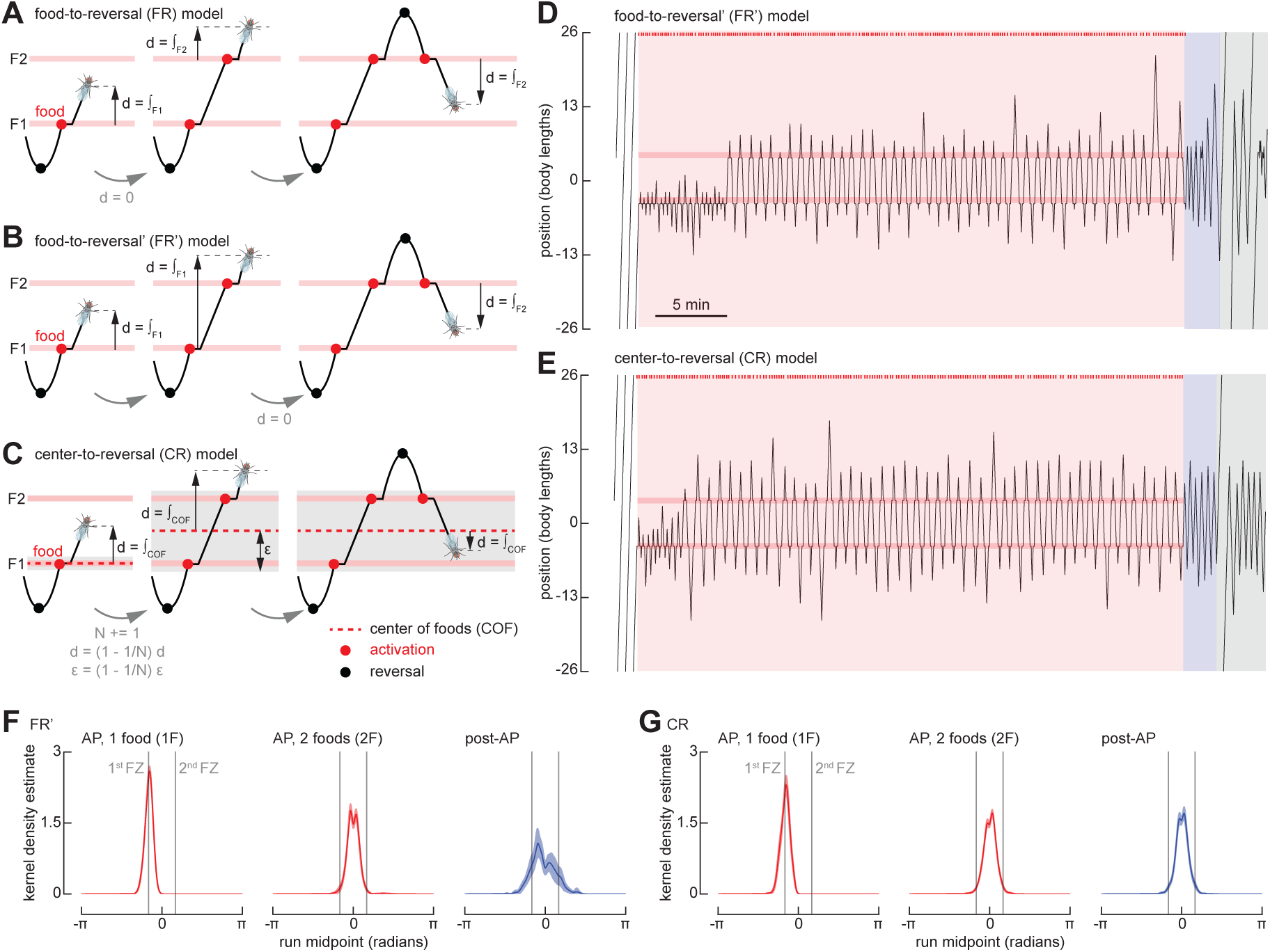
Two modified versions of the FR model recapitulate *Drosophila* search behavior around multiple fictive food sites. **(A)** Schematic, showing features of food-to-reversal (FR) model. The virtual fly resets its integrator at each new food that it encounters. **(B)** As in (A) for food-to-reversal (FR’) model. The virtual fly resets its integrator at the first food encountered after each reversal. **(C)** As in (A) for center-to-reversal (CR) model. The virtual fly resets its integrator at the center of all the food locations it encounters during a run. **(D)** As in Figure 4A, for a simulation using the FR’ model. **(E)** As in Figure 4A, for a simulation using the CR model. See also Video S4. **(F)** As in Figure 4B, for simulations using the FR’ model. The first 300 simulations in which the virtual fly found both food sites are included. (N = 300). **(G)** As in Figure 4B, for simulations using the CR model. The first 300 simulations in which the virtual fly found both food sites are included. (N = 300).

An alternate, the center-to-reversal (CR) model (Figure 5C, Video S4), is based on the notion that the fly can compute and store the coordinates of a location that is at the center of a cluster of food sites—the center of foods (COF); it is at this location where the path integrators begin to accumulate. When only one food site is present, the COF coincides with the food site and the model integrates from that location, as in the FR and FR’ models. However, when the fly encounters a second food site, it shifts the origin of its search to a location between the two food sites. To distinguish new food sites from previously encountered sites, the simulated fly also calculates the rough size of the food patch, ε. If the fly encounters a food site when its integrator exceeds ε, both the integrators and ε are updated such that the origin is always at the COF. The model implements a simple algorithm to place the COF at the center of mass of all known food sites and to calculate ε as the distance from the COF to the outermost food site. We are not inferring that this is a biologically plausible mechanism by which such calculations might be implemented—indeed, many simple mechanisms are possible to maintain an estimate for the center of the food patch. The important distinction is that, in the CR model, the fly can estimate the center of the food sites it has encountered, and that this calculation requires some short-term memory; its behavior cannot be explained by a reflexive action to the last food site visited. Despite the fact that the model parameters were determined via a grid search on the one food configuration dataset, both the FR’ and CR models do a reasonable job of recapitulating the flies’ behavior in a two food geometry, in that KDEs of the run midpoint distributions are unimodal during both the AP and post-AP periods (Figure 5F-G).

To test between the FR’ and CR models, we modified the arena to feature three food zones, spaced 4.5 body lengths apart (Figure 6A). We changed our optogenetic protocol such that, at the end of the AP, we disabled all but one of the food zones, which remained active for just one single additional visit (Figure 6B). The FR’ and CR models make very different predictions under these conditions. In the FR’ model, the position of the expected run midpoint during the post-AP is strongly dependent upon the position of the last food site encountered, whereas this is not the case with the CR model. As shown in Figure 6, the run midpoint KDEs measured from real flies are indistinguishable in the three experimental conditions (final food at top, middle, or bottom position), and were centered at a point corresponding to the middle food site, which matches the prediction of the CR model. These results support the basic assumptions of the CR model, which is that the fly can somehow retain and update a memory of the center of the food patch over time as it zig-zags back and forth across the individual food sites, and that this memory is not dependent on the location of the last active food site it encountered. The CR model also successfully recapitulates flies’ behavior in reinitiating the search after completing one or more circles around the arena (Figure S4 and Video S5).

**Figure 6.**
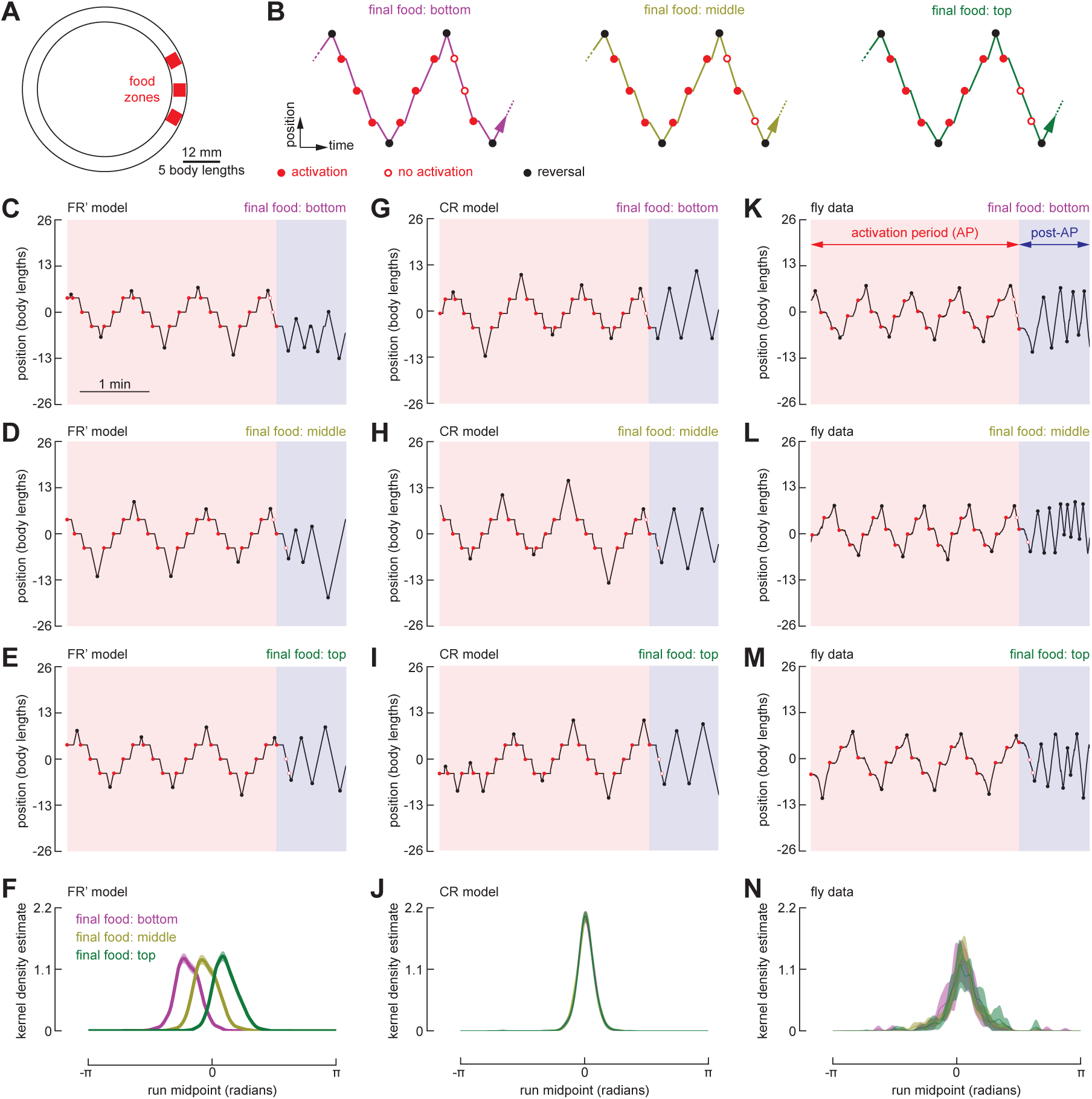
Flies reset their path integrator at the center of a cluster of fictive food sites. **(A)** Schematic of the annular arena with three food zones, spaced 4.5 body lengths apart. **(B)** Schematic of the experimental paradigm. At the conclusion of the AP, two of the food zones were disabled while one food zone remained capable of providing an additional single optogenetic pulse. For each trial, the final operational food zone was designated to be either the bottom, middle, or top. For trials where the final 3 or more runs during the AP spanned all three food zones, we calculated the KDE of the run midpoint during the post-AP. **(C-E)** Example trajectories of the FR’ model fly searching across three food zones, in which the final food stimulus is in either the bottom (C), middle (D), or top (E) position. **(F)** Normalized KDE of the run midpoint in the post-AP period for FR’ model. (N = 300). **(G-J)** As in (C-F), for simulations using the CR model. (N = 300). **(K-N)** As in (C-F), for fly data. (N = 45 flies, n = 166 trials).

## DISCUSSION

In this paper, we developed a novel assay to study path integration during foraging in *Drosophila*, which provides quantitative insight into a behavior that is more difficult to analyze in a simple open field arena. We induced local search in a ring-shaped channel by optogenetically stimulating sugar receptors whenever the fly occupied one or more arbitrarily defined food zones. Local searches in this arena manifested as a persistent zig-zag pattern in which flies iteratively walked away from and back to the food zone (Figures 1B-C)—a pattern that persisted even after the optogenetic stimulation was no longer provided. After the optogenetic activation was disabled, the flies continued to walk back and forth around the site where they had experienced food. While we cannot directly measure what is going on in the brain of a freely walking animal, we derived a convenient proxy for the flies’ spatial memory by constructing a kernel density estimate (KDE) based on the midpoints of the back and forth runs. An examination of the KDE in any given period provides a clear measure of where in the arena a fly was centering its search. In experiments in which the flies were presented with just one food site, this KDE function remained centered at the single food location after the activation protocol was switched off (Figure 1M). In experiments in which we presented the fly with two food sites, the peak of the KDE moved to a point midway between two food locations (Figure 4B). These data suggest that the flies adjust their integrator such that the zero location is midway between two food sites, even though there is no sensory stimulus associated with that location. This interpretation is consistent with the agent-based modeling we performed, in that only the CR model, which maintains the zero location of both of the path integrators used to keep track of the simulated fly’s position at a location in between food sites, could recapitulate the behaviors we observed (Figures 5 and 6). This apparent ability to center the idiothetic integrator at a location not directly linked to a sensory stimulus identifies a hitherto unknown capability of the navigation system in *Drosophila*.

In nearly every fly we examined, individuals did not give up local search gradually, but rather executed what we termed a departure run, which was substantially longer than all previous runs in the post-AP (Figures 1J-L). This suggests that the behavioral state of the fly may change over the time course of several minutes after it stops receiving a food stimulus. Nevertheless, in a smaller circular arena, we found that some flies reinitiated a local search at the food site after traveling one or more times around the circumference of the chamber (Figure 2), indicating that they retained a spatial memory of the food location and presumably had not changed in physiological state to the point that they had given up their search for the food. Whereas the short back-and-forth motions of the fly around the food site during the AP and post-AP might be explained by simple one-dimensional odometry, the flies’ ability to recognize the food site after circling the arena cannot. Such a feat instead requires that flies integrate azimuthal heading information with odometry to perform true two-dimensional path integration. Because our experiments were conducted in the absence of visual or chemical cues, and the results were robust to manipulations such as switching the location of the food stimulus on a trial by trial basis (Figure 2), we presume flies measure their translation in the arena via idiothetic self-motion cues, such as proprioception or efference copy of motor commands^9^. The obvious candidate locus for the computations associated with our hypotheses is the Central Complex (CX), a set of unpaired neuropils in the core of the insect brain^8, 9, 32^. Recently, work on CX neuroanatomy^33, 34^ and physiology^35–37^ has characterized a network of neurons that encode compass-heading, leading to models wherein these circuits provide the angular heading information required for path integration. Although mechanisms by which odometric information is encoded and read out by the CX have been proposed^32, 38–40^, none have yet been explicitly tested via genetic or physiological manipulation of behavior.

Our experimental setup in which the flies are restricted to a narrow, dark channel is similar in some regards to the heat box paradigm developed by Wustmann and colleagues^41^. In that apparatus, flies avoid one half of the chamber that is heated to an aversive temperature and retain the avoidance after the temperature stimulus has been removed. This avoidance has been interpreted to depend on path integration, and clever use of this assay in combination with genetic tools has implicated populations of serotonin cells in the central brain as being critical for place memory^42^. Ofstad and coworkers^43^ presented another aversive spatial memory paradigm, based on prior work in cockroaches^44^, in which flies learn the position of a cool patch within a hot circular arena using visual cues. Genetic silencing experiments using that paradigm implicated the ellipsoid body (EB) as being critical for place memory. The implication of the EB in this place memory paradigm makes sense in the light of more recent work demonstrating the existence of a compass cell network within that structure. The paradigm described in this paper is different in that the stimulus (optogenetic activation of sugar receptors) that drives the formation of spatial memory is attractive and relies on the behavioral modules employed during foraging behavior rather than those associated with heat avoidance. However, the difference in the representation of attraction and aversion in the CX remains unclear, as does the driving force for resetting the path integrator to a target location. Our assay also provides precise control over the presentation of such stimuli, enabling for the interrogation of the time course of the formation of spatial memory. To systematically test the time course, one might dynamically control the two food sites to see, for example, how many experiences with a second food site are required for the fly to exhibit a two-food search modality in the post-AP. Anecdotally, we observed two-food searches in the post-AP after very few experiences with the second food site or after periods in which the fly has reverted to a one-food search in the AP (Figure S1), but a formal analysis of these temporal dynamics was beyond the scope of the current study.

Unlike a prior computational model of path integration by Stone and coworkers^32^, our agent-based, state-transition models are not constrained by neural architecture and physiology. However, their output may be directly compared to those of actual experiments, allowing us to probe whether the flies’ behavior might require certain cognitive capabilities. Further, the models were not only testable but endowed with predictive power, such that the generation of models satisfying the one food dataset inspired experiments to distinguish between these models. With much effort, we tried to build a model (i.e., our FR’ model) that could explain a fly’s ability to center its search around a location in the middle of two food sites based only on path integrators that re-zero at the location of food. However, the FR’ model unambiguously failed to predict the results of our three food experiments, which instead indicate that the flies’ spatial memory is not determined by the location of the last active food site it encountered (Figures 5C and 6). As suggested by the success of the CR model, flies appear to accumulate experience while foraging to develop some internal sense of the food patch’s center.

While the CR model successfully recapitulates all aspects of the flies’ behavior, its neural implementation is not clear. Performing a search around a central location in a cluster of food sites, as in the two- and three-food experiments, can be explained by at least two mechanisms. First, given the ability of honeybees to count landmarks^45^ and trapline across multiple food sites^46^, the flies could be treating multiple food sites as individual loci within a larger food patch, and adapt their search to cover the entire patch. This mechanism could require that flies have memory of multiple food locations, but a more parsimonious explanation may be that only the central place is stored and each food experience reinforces or updates the place memory of the central location. Alternatively, the food sites could be interpreted as a single food source with high uncertainty in its location; in accordance with optimal foraging strategies, the maximal search radius of the fly would increase with uncertainty as the square root of the time since the last food encounter^6, 17, 47^. This hypothesis is consistent with our observation for the increase in run lengths at the beginning of the post-AP, but not with the plateau seen later in the post-AP (Figure 1H). Furthermore, whereas zeroing the path integrators in the models refers to resetting the integrated value to zero, the corresponding biological mechanism is likely more complex. The second mechanism suggested here—that the food sites are interpreted as a single location with high uncertainty— could be explained by an incomplete zeroing of the path integrators upon discovery of the second food site. If memory decay is mediated by, for example, time spent feeding, then residual memory might decay insufficiently to center the search at the new food site and instead cause the fly to search over a central location. Modeling the possibilities separately (as opposed to a general central place model presented here) combined with experiments (e.g., presenting stimuli of differing reward strengths or durations) could help distinguish between these possibilities.

In all the experiments reported in this paper, we constrained flies to an annular arena, simplifying the complex, tortuous paths of flies performing local search in an unconstrained environment to sequences of runs separated by changes in direction (i.e., reversals). The convenient linearization of the behavior allowed us to focus much of our analyses and modeling on the reversals, events for which there is no direct equivalent in open field arenas. Even so, reversals likely correspond to behaviors seen in the local searches of unconstrained flies. For example, in an open arena, flies typically walk in straight lines interspersed with discrete changes in direction, which can be as small as a few degrees or as large as 360°^27, 28^. Reversals in the annular arena could correspond to the fly attempting to turn by an angle above a certain threshold. Another possible explanation is that the flies accumulate their changes in direction and reverse course when the accumulated attempted turn angle reaches a threshold. A third possibility is that the reversals are, in fact, unrelated to turn angles in open arenas; rather, the fly executes a reversal when it has reached the maximal distance away from the food site it is willing to venture within the channel.

In sum, we have developed high-throughput assays to quantitatively measure path integration in *Drosophila*, with the ability to quantify spatial memory by measuring the midpoint locations of the runs executed by the animals. The results support the hypothesis that flies employ path integration during foraging behavior, and that they can use this system to center their search at a featureless location situated in between a cluster of food sites. Future studies might employ these assays, in combination with genetic manipulation of neural activity, to further unravel the neural mechanisms of path integration.

## ACKNOWLEDGMENTS

We would like to thank all members of our lab for helpful discussions. Will Dickson provided essential help with programming and construction of our experimental set-up. Research reported in this publication was supported by the National Institute of Neurological Disorders and Stroke of the National Institutes of Health under Award U19NS104655.

## AUTHOR CONTRIBUTIONS

AHB conducted all experiments, under the supervision of MHD. EHP developed the state-based models of behavior. AHB, RAC and EHP analyzed data and prepared all figures. AHB, RAC, EHP and MHD wrote the paper.

## EXPERIMENTAL MODEL AND SUBJECT DETAILS

We conducted all experiments using 3-to-6-day-old female *Drosophila melanogaster* reared in darkness at 22°C. We reared the flies on standard cornmeal fly food containing 0.2 mM all trans-Retinal (ATR) (Sigma-Aldrich) and transferred flies 0-2 days after eclosion onto standard cornmeal fly food with 0.4 mM ATR. We supplemented the standard cornmeal food with additional yeast. We obtained the flies by crossing *Gr5a-Gal4* male flies with *UAS-CsChrimson* female virgin flies. Prior to experiments, we wet-starved flies by housing them for 24-40 hours in a vial supplied with a tissue (KimTech, Kimberly-Clark) containing 1 mL of distilled water with 800µM ATR, and dry-starved flies for up to 150 minutes, including a 45-to-90-minute acclimation period in the experimental arena.

## METHODS DETAILS

### Behavioral experiments with walking flies

We conducted all experiments in a 40 mm-diameter annular arena, except for experiments in Figure 2, where we used a 20 mm-diameter annular arena to increase the likelihood that flies would complete full revolutions during the post-return period. We constructed the arenas from layers of acrylic with insertable acrylic discs to create the annular channel (4 mm wide and 1.5 mm high). The width of the channel provided sufficient space for flies to walk forward, backward, or turn around at any point in the arena. The channel’s low height encouraged the fly to walk either on the floor or the ceiling, rather than the walls of the channel. An upward-directed, custom-made array of 850 nm LEDs, covered by a translucent acrylic panel, was situated 12 cm beneath the arena to provide backlighting for a top-mounted camera (blackfly, FLIR) recording at 30 frames per second. For optogenetic stimulation, we positioned upward directed, 628 nm LEDs (CP41B-RHS, Cree, Inc.) at the center of each food zone, 8.5 mm beneath the arena floor. We covered the chamber lid with a 3 mm thick long-pass acrylic filter (color 3143, ePlastics). The chamber floor was transparent to allow for optogenetic stimulation, and a filter (#3000 Tough Rolux, Rosco Cinegel) was situated beneath the chamber to diffuse the red light used for stimulation, resulting in ∼300 W of illumination at the arena floor. The camera, fly chamber, optogenetic lighting panel, and background lighting panel was held within a rigid aluminum frame (80/20) covered with black acrylic to block any external light. We tracked the 2D position of the fly in real time using a python-based machine vision system built on the Robot Operating System (http://florisvb.github.io/multi_tracker). We customized the tracking software to implement closed-loop control of optogenetic stimulation via an LED controller (Arduino Nano). During our initial experiments, we cleaned the behavioral chamber with 100% ethanol after each trial and allowed it to dry before reuse. However, because ethanol causes cracks in the acrylic parts, we stopped using ethanol and instead cleaned the chambers with compressed air. We did not observe any difference in fly behavior between the two cleaning methods (data not shown).

For each experiment, we aspirated a single fly into the behavioral chamber, allowing it to acclimate for 45-90 minutes. The final minutes of this acclimation period correspond to the baseline period in our analyses. Following acclimation, experiments consisted of a specified time-course of activation periods (APs) and post activation-periods (post-APs). During APs, the LED beneath each food zone was turned on for 1 s whenever the centroid of the fly occupied its virtual perimeter (=2.6 BL or 1.3 BL for the small arena experiment in Figure 2). Because optogenetic activation of sugar sensors inhibits locomotion^28^, each 1 s pulse was followed by a 15 s refractory period during which the LED remained off, regardless of the fly’s position. During the baseline period and post-APs, food zones were not operational such that flies could not receive optogenetic activation.

For experiments with multiple APs (as in Figure 1C), each AP and subsequent post-AP was treated as a single trial. For all experiments with a 40-min AP (e.g., Figure 1B), data were discarded if the fly moved less than 10 cumulative body lengths during the first 20 minutes of the AP (N = 3 flies discarded). For all trial-based experiments (e.g., Figure 1C) APs, trials were discarded if the fly moved less than 10 cumulative body lengths during the AP (n = 22 trials discarded).

For experiments in Figure 2, to discard the possibility that flies were able to find food zones by sensing temperature gradients generated by the LEDs, one of the control zones was outfitted with an LED. When food stimuli were presented, the LEDs at both food zones as well as this control zone were turned on.

During experiments in Figure 6, we encouraged flies to expand their search to span all three food zones by using an altered protocol during the AP in which we disabled each food zone after it was encountered by the fly for the first time; after the fly had encountered all three food zones, all the food zones became operational and remained operational for the remainder of the AP.

### Agent-based models without odometric integration

The random sampling and Lévy flight models simulated post-AP search by drawing from natural statistics derived from fly search trajectories. For the random sampling model, run lengths were randomly sampled with replacement from the fly post-AP run lengths in Figure 1J (excluding the departure run). For the Lévy flight model, a Lévy distribution was fit to the same data using the function stats.levy.fit() from SciPy, and run lengths were drawn randomly from the resulting distribution.

### Agent-based models featuring odometric integration

The food-to-reversal (FR), food-to-reversal’ (FR’), and center-to-reversal (CR) integration models are graphically described by the state transition diagrams in Figure S3. The fly is simulated as a point-mass within a virtual environment (Figure S3A) consisting of an annular channel, 52 body lengths (BL) in circumference. The environment includes one or more food zones (1 BL in length) at specified locations along the linear channel. Similar to our experiments with real flies, whenever the simulated fly enters a food zone in the simulated environment, it receives a 1 s food stimulus, followed by an 8 s refractory period during which the simulated fly cannot receive a food stimulus; whereas the refractory period is briefer in the simulations than in experiments with real flies, comparisons of temporal aspects of the two systems are somewhat arbitrary because the walking speed of simulated flies is defined artificially. When walking, the fly moves 1 BL per time step and corresponds to 0.5 s (i.e., the simulated fly walks at 2 BL s^-1^).

The fly is initialized at the 0 BL position, in the counterclockwise orientation, and the simulated environment (Figure S3A) is initialized in the food-off state. The fly’s integrators are initialized with a value of 0 and the fly’s target run length value, *r_t_*, is initialized to 0. When the environment is in the food-off state, at each time step, the system checks whether the current time is during the AP, whether the current time is not during a refractory period (i.e., whether the time since the last food stimulus exceeds the duration of the refractory period), and whether the fly occupies a food zone; if all of these conditions are satisfied, the food stimulus is turned on and the environment enters the food-on state. When the environment is in the food-on state, at each time step, if the food stimulus has been on for the duration of the specified stimulus duration (1 time step), the food stimulus is turned off and the environment returns to the food-off state. The state transition diagram described in Figure S3A—with varying food zone positions as well as varying baseline, AP, and post-AP durations—is used to simulate the environment in all the models (FR, FR’, and CR).

### Food-to-reversal integration model

In the food-to-reversal (FR) model (Figure S3B), the simulated fly is able to measure walking distance using two integrators—one integrator measuring displacement along the North-South axis, I_NS_, and a second measuring displacement along the East-West axis, I_EW_. These directions are defined within the simulated fly’s reference frame, rather than a global reference frame in the simulated environment; that is, while the environment imposes a global reference frame, the simulated fly constructs its own map of space agnostic to the absolute directions an observer may impose (e.g., the simulated fly may assign North to the direction an observer would call West). The simulated fly is able to store and retrieve its previous action—either a reversal or an eating event. The simulated fly is initialized in the global search mode in the walking state, where it moves forward 1 BL at every time step. When the fly receives a food stimulus, it transitions to the eating state in the local search mode. The fly remains in the local search mode for the remainder of the simulation. While the fly continues to receive the food stimulus, it remains in the eating state. Upon delivery of a food stimulus, the simulation is advanced 10 time steps, during which the fly remains stationary, which mimics the locomotory pause induced by activation of sugar-sensing neurons in *Drosophila*. While flies exhibit more complex behaviors when walking in the absence of food stimuli (e.g., pausing, changes in walking speed) and in response to food stimuli (e.g., proboscis extension), the models aimed to reduce the behavior to its simplest form to exclusively interrogate the bounds of integration, so these more complex modalities were ignored.

Upon the termination of the food stimulus in the FR model, the fly selects a new target run length, *r_t_*, by drawing a value from C_f_, the distribution of food-induced run lengths. As described in a subsequent section, it is not possible to directly observe C_f_ in data from real flies, and we therefore derived this distribution via an optimization procedure that fits our model to data. Upon termination of the food stimulus in the FR model, the integrators are both set to zero, such that the food site serves as the origin of the fly’s search. This behavior is in accordance with traditional PI models, wherein the integrators are zeroed at the origin of search and a maximal excursion distance is selected.

Having responded to the food stimulus, the simulated fly sets its previous action to an eating event and transitions to a walking state. At each time step while in the walking state, the fly moves forward 1 BL, and the integrators are incremented by decomposing the step into orthogonal components along the North-South and East-West axes using trigonometric functions of the fly’s heading direction, θ_h_. The fly receives its heading direction from the environment at every time step rather than including a mechanism in the model for the fly to determine its heading based on idiothetic or allothetic cues.

The fly recalls whether a full revolution has been made since last passing the food site to reinitiate the search after full revolutions. In these cases, a run length has been selected which exceeds the maximum possible integrator value when the fly is constrained to the circle, so the fly will never perform a reversal and, in the post-AP, never select a new run length. When the absolute value of the heading angle exceeds 3 rad (172°), the fly recognizes that a full revolution has occurred. Alternatively, if the absolute value of the heading angle falls below 0.15 rad (8.6°) and a full revolution has occurred, the fly has returned to the food site and, accordingly, chooses a new run length, zeroes its integrators, and sets the full revolutions variable to False. In essence, this feature of the model enables the simulated fly to recognize its return to the food site despite not having made a reversal and to choose a new run length and resume search accordingly.

The fly continues walking until the Euclidean norm of the integrators equals or exceeds its current target run length, *r_t_*. At this point, a new target run length is selected based on the fly’s previous action. If the previous action was an eating event, *r_t_* is defined to be the sum of the value of the Euclidean norm of the integrators and a value drawn from C_f_; this ensures that the search stays centered over the food zone. On the other hand, if the previous action was a reversal, *r_t_* is defined to be the sum of the Euclidean norm of the integrators and a value drawn from C_Δ_, the distribution of the differences in lengths between consecutive runs. As described in a subsequent section, we determine C_Δ_ via an optimization procedure that fits our model to data from real flies. After the selection of a new target run length, walking direction is reversed and the integrators are zeroed. The fly remains in the walking state and returns to the eating state if it receives a food stimulus. This is in accordance with traditional path integration models, wherein the agent only searches within a certain distance from the origin; constrained to a one-dimensional environment, the agent executes a reversal when this limit is reached.

### Food-to-reversal’ integration model

In the food-to-reversal’ (FR’) model (Figure S3C), the simulated fly measures walking distance using four integrators—one integrator for displacement in each direction: North, South, East, and West (I_N_, I_S_, I_E_, and I_W,_ respectively). The FR’ model would not function with only two orthogonal integrators because the simulated fly must keep track of two distances to accomplish local search during the post-AP in environments with multiple foods: the distance between the foods and the distance walked since the last reversal. In a two-food configuration, for example, immediately preceding a reversal, the integrators in the previous direction of travel will store the distance to the further food site and the opposing integrators will both be zero. After the reversal, the integrators storing the distance to food will begin to decrement and the others will increment. Upon encountering the closest food site, the first set of integrators will then exactly equal the distance between the food sites, whereas the second set of integrators will recall the distance from the reversal to the current location. Thus, the model measures the distance from the reversal, as did the FR model, while still recalling the distance between food sites in multiple food configurations. Like the FR model, however, feeding sites remain the locations where all integrators re-zero, but with differences described more fully below.

Upon the termination of the food stimulus in the FR’ model, the fly selects a new target run length, *r_t_*, based on the fly’s most recent previous action. If the fly’s most recent action was a reversal, then *r_t_* is defined to be the sum of I_o_, a value computed as the Euclidean norm of the two integrators opposing the current direction of travel, and a value drawn from C_f_, all integrators are set to zero, and θ_h_*, the food site heading angle, is set to the current heading angle. This course of action represents the fly responding to having received its first food stimulus since performing a reversal; in the FR’ model, the first food stimulus after a reversal is treated as the origin of search, so the integrators are zeroed and a run length is selected accordingly. Similarly, if the most recent action was a full revolution, *r_t_* is a value drawn from C_f_, the integrators are zeroed, and the food site heading angle is set to the current heading angle; this course of action represents the fly encountering a food site after performing a full revolution around the circle. On the other hand, if the fly’s most recent action was an eating event (the only possible action other than a reversal or a full revolution) and the difference between the current target run length and the value of I_d_, a value computed as the Euclidean norm of the two integrators aligned with the current direction of travel, is below 1 BL, then the new target run length, *r_t_*, is defined to be the sum of the value of whichever integrator is highest and a value drawn from C_f_; this course of action represents the simulated fly interpreting the food stimulus as a new food location and extending its run length to expand its local search to encompass the new food in addition to the prior food(s). Finally, if none of the conditions holds, the fly does not select a new target run length; this course of action represents the fly encountering a food site that has been previously experienced and which the search has already been expanded to encompass. In sum, the fly sets the first food site after a reversal as the origin of its search, does not change the origin in response to additional food sites on that run, and only extends the run length if a previously unexperienced food site is encountered.

Having responded to the food stimulus, the simulated fly sets its previous action to an eating event and transitions to a walking state. At each time step while in the walking state, the fly moves forward 1 BL, and the integrators are incremented accordingly with a minimum value of zero. The mechanism for identifying and responding to full revolutions is similar to that of the FR model; here, however, the fly uses the difference between the heading angle and the food site heading angle such that a full revolution is only identified when the fly’s position diametrically opposes the position of the food site.

As in the FR model, the fly executes a reversal when it reaches the target run length, the maximal distance from the origin of search it is willing to venture. Algorithmically, this distance is identified as I_d_ equaling or exceeding *r_t_*. At this point, a new target run length, *r_t_*, is selected based on the fly’s previous action. If the previous action was an eating event, *r_t_* is defined to be the sum of the value of I_d_ and a value drawn from C_f_; this ensures that the search stays centered over the food zone(s). On the other hand, if the previous action was a reversal, *r_t_* is defined to be the sum of I_d_ and a value drawn from C_Δ_. After the selection of a new target run length, walking direction is reversed. As the integrators are strictly positive, the integrators in the direction of travel following a reversal are always zero, so a formal zeroing step is not required. The fly remains in the walking state and returns to the eating state if it receives a food stimulus.

### Center-to-reversal integration model

In the center-to-reversal (CR) model (Figure S3D), the simulated fly is able to measure walking distance using just two integrators, I_NS_ and I_EW_. Furthermore, the simulated fly is able to store and retrieve its previous action—either a reversal or an eating event. As in the FR and FR’ models, the fly is initialized in the global search mode in the walking state and transitions to the local search mode, eating state upon receiving a food stimulus. The fly remains in the local search mode for the remainder of the simulation.

After receiving a food stimulus, the fly stays in the eating state until the termination of the food stimulus, at which time it transitions to the walking state. If the fly’s previous action was eating, a new run length is selected as the sum of a value drawn from C_f_ and ε, a term that represents its rough estimate of the expanse of the food region encompassing multiple food sites. If the Euclidean norm of the integrators exceeds ε, the fly recognizes that it has encountered a new food site and adjusts its integrators such that the origin of its search is at the center of mass of the food sites. Additionally, ε is adjusted to exceed the distance from the center of mass to the outermost food site. To accomplish this, it increments N, the total number of food sites present, by one, and updates the integrators and origin heading angle by multiplying them by a factor of 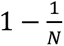. This effectively shifts the center of mass by the current integrator value along its axis divided by the number of food sites, such that the center location is always the average location of all known food sites. The expanse, ε, is also updated by multiplication of the Euclidean norm of the integrators by the same factor, and a further multiplication by 1.5 to ensure the expanse encompasses all food sites regardless of numerical errors.

Having responded to the food stimulus, the simulated fly sets its previous action to an eating event and transitions to the walking state. While walking, the integrators are incremented via orthogonal components of the step length as in the FR model. When the Euclidean norm of the integrators falls below one, a new target run length is selected such that if the fly performs a full revolution, it reinitiates the search at the center of the food site.

As in the previous models, when the Euclidean norm of the integrators exceeds the target run length, the fly performs a reversal. The new target run length is defined to be the sum of the expanse and a value drawn from C_f_, the direction is reversed, and the fly returns to the walking state. The fly remains in the walking state and returns to the eating state if it receives a food stimulus.

### Run length distributions for odometric integration models

In all three models, the simulated fly selects a new target run length following a food stimulus by sampling from the food-induced run length distribution 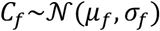. To select a new target run length following a reversal, the FR and FR’ models sample from 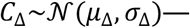—the distribution of the difference in length between consecutive runs—whereas the CR model always draws from *C_f_* when selecting a new run length, regardless of previous actions. Whereas the sampled distributions are analogous to the observable statistics of *Drosophila* local search, they cannot be derived from fly data, because we cannot directly measure target run length in a real fly. For example, when a fly encounters a new food location and continues walking several body lengths before performing a reversal, the resulting total run length might be the sum of the original target run length (selected prior to encountering the new food) and an additional run length induced by the new food stimulus; therefore, we cannot directly measure the true value of either component of the fly’s algorithm. Instead, we determined the distribution parameters by performing a grid search over the parameter space to minimize a cost function (Equation 1). At each point in the grid search, the model was run 250 times in the one food configuration. The cost function was designed to minimize the differences between the statistics of *Drosophila* local search and those of the given model. Given *N* parameters we sought to match between the data and simulations, we fit an appropriate distribution (e.g., inverse gaussian) to the *i^th^* parameter to get the distribution 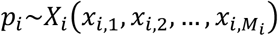, such that the distribution is governed by *M_i_* values. We fit the distribution to both the data and the simulations, yielding 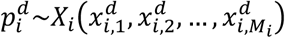 for the data and 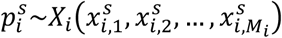 for the simulations (where *d* and *s* denote ‘desired’ and ‘simulated’, respectively). We then calculated the total cost across all parameters, normalizing for the number of values governing each distribution:

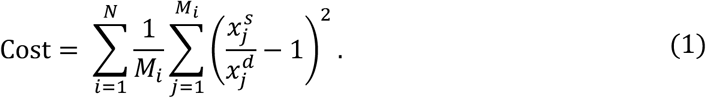

The relevant parameters we sought to match between the data and simulations were the excursion distances 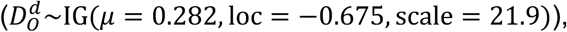, the run lengths in the post-AP 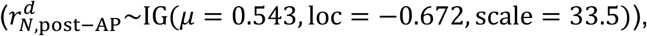 and the locations of run midpoints 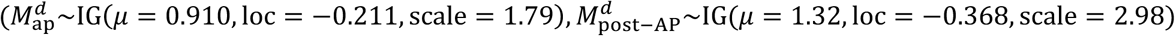. These values were derived using SciPy’s stats.invgauss.fit() function. The distributions used in the FR model were 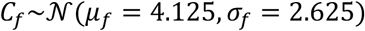 and 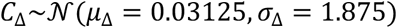. The distributions used in the FR’ model were 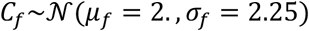 and 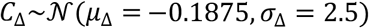. The distributions used in the CR model were 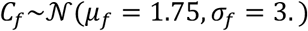. The final cost for the FR model was ∼76.1, the final cost for the FR’ model was 2.14, and the final cost for the CR model was 3.06.

### Behavioral analysis of walking flies

The dataset for each experiment consisted of an array of X and Y coordinates representing the 2D positions of the fly, as well as an array of LED states (on or off) for each food zone. Data were sampled at ∼30 Hz. We converted the positional coordinate of the fly to an angular position in the ring-shaped arena and treated the fly as a point mass along the circumference of the arena. The beginning of each AP was defined as the first food stimulus, and the end of each AP (and the beginning of the subsequent post-AP) was defined as the final food stimulus. To process data, we discarded occasional frames where the fly was either not tracked, where a second object was tracked in addition to the fly (e.g., fly poop), or where the tracked position jumped more than 3 mm within two consecutive frames (e.g., due to sporadic tracking of another object). Because the position of food zones varied slightly due to variations in the fabrication and assembly of arenas, we defined the center of each food zone for each experiment as the midpoint between the extrema of fly locations at food stimulus events associated with the food zone.

## QUANTIFCATION AND STATISTICAL ANALYSIS

We generated all figures using the python library matplotlib. Throughout the paper, we calculated the 95% confidence intervals using built-in SciPy statistical functions to compute the standard error of the mean and the Student’s t-distribution. For the statistical significance analysis, we used distributions of mean values generated by 2000 bootstrap iterations. For datasets with overlapping discretized ranges, we plotted datapoints in a random order for presentation purposes.

## Supplemental Video Legends

**Video S1. Animations of *Drosophila* local search behavior, related to** **Figure 1**.

Each animation depicts fly position (black circle) during one trial from the experiment plotted in Figure 1C. A motion trail depicts fly position during the previous 5 seconds. Optogenetic activation events are indicated by a red circle to the left of the food zone. The total elapsed time and experimental period are indicated. Playback is at 8x speed.

**Video S2. Animations of *Drosophila* local search behavior in a small annular arena, related to** **Figure 2**.

Each animation depicts fly position during one trial from the experiment plotted in Figure 2B. A motion trail depicts fly position during the previous 5 seconds. Plotting conventions as in Video S1. Playback is at 8x speed.

**Video S3. Animation of *Drosophila* local search around two food zones, related to** **Figure 4**.

The animation depicts fly position during the experiment plotted in Figure 4A. A motion trail depicts fly position during the previous 12 seconds. Plotting conventions as in Video S1. Playback is at 20x speed.

**Video S4. Animations of simulated data from CR model of local search around two food zones, related to** **Figure 5**.

Each animation depicts virtual fly position during the AP and post-AP from CR model simulations. A motion trail depicts fly position during the previous 10 seconds. Plotting conventions as in Video S1. Playback is at 18x speed.

**Video S5. Animations of simulated data from CR model in a small annular arena, related to** **Figure 2**.

Each animation depicts virtual fly position during the post-AP and post-return from CR model simulations. A motion trail depicts fly position during the previous 10 seconds. Plotting conventions as in Video S1. Playback is at 16x speed.

---

**Figure S1.**
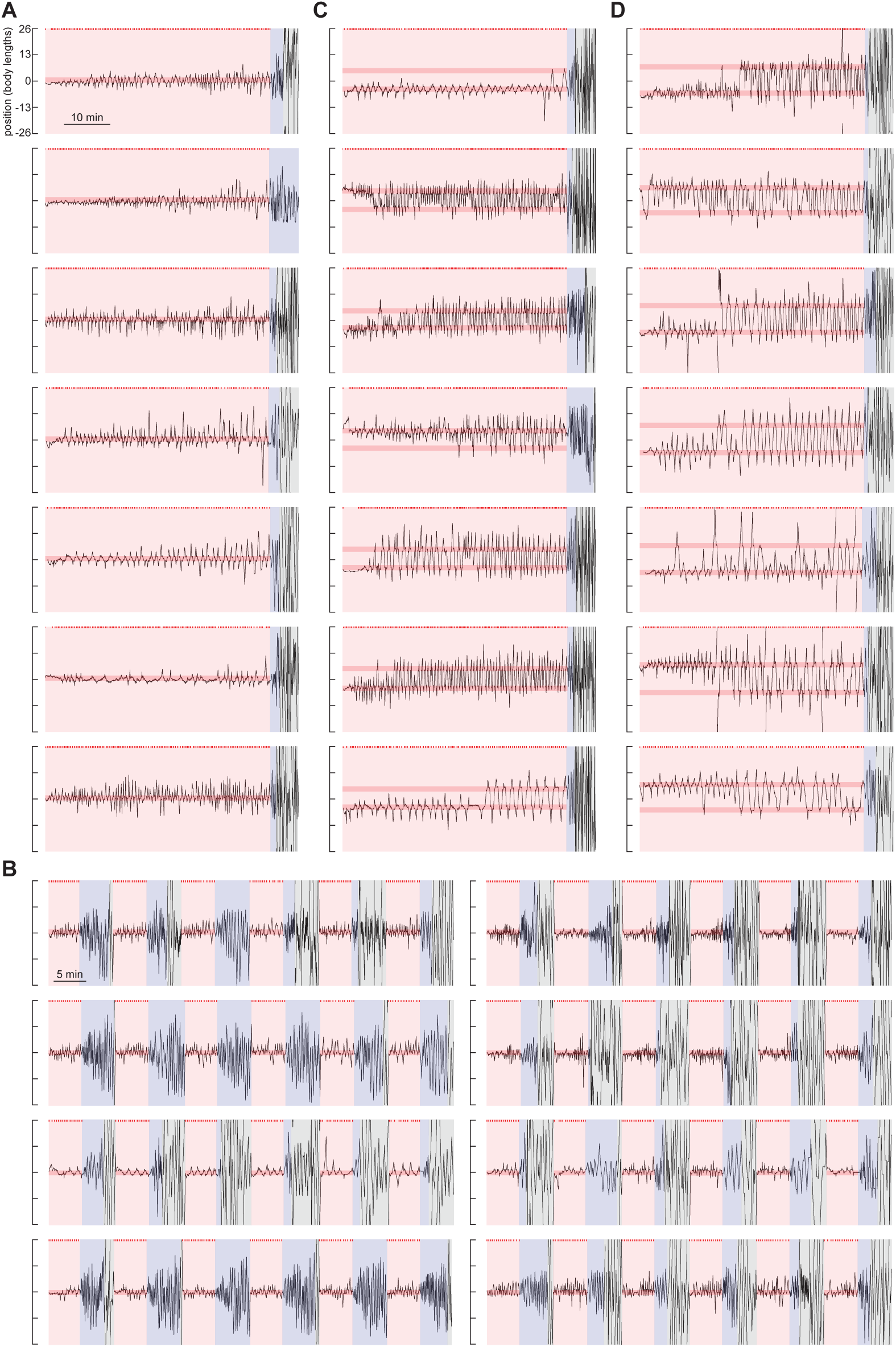
Sample trajectories of local search around one or two fictive food sites, related to Figures 1 and 4. **(A)** Seven representative example trajectories for the AP and post-AP from single food 40-min AP experiments. Plotting conventions are the same as Figure 1B. **(B)** Eight representative example trajectories for the AP and post-AP from single food trial-based experiments. Plotting conventions are the same as Figure 1C. **(C)** Seven representative example trajectories for the AP and post-AP from two-food 40-min AP experiments. The two food zones are spaced 9 body lengths (BL) apart. Plotting conventions are the same as Figures 1B and 4A. **(D)** Seven representative example trajectories for the AP and post-AP from two-food 40-min AP experiments. The two food zones are spaced 13 body lengths (BL) apart. Plotting conventions are the same as Figures 1B and 4A.

**Figure S2.**
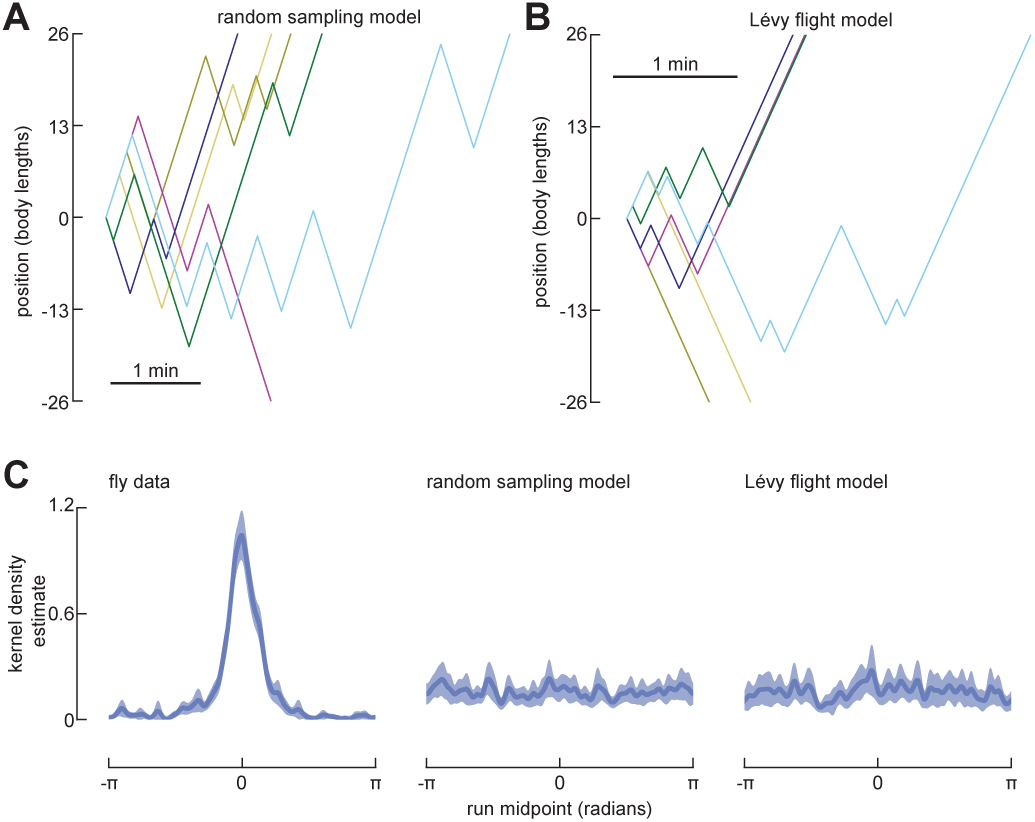
Memory-less models cannot recapitulate *Drosophila* local search, related to Figure 3. **(A)** Six representative example trajectories from simulations for which run lengths were randomly drawn from the distribution of run lengths in Figure 1J (excluding the departure runs). Trajectories begin at the 0 position and are terminated when the simulated fly reaches 26 body lengths from the point of origin. **(B)** As in (A) from simulations for which run lengths were drawn from a Lévy distribution fit to the distribution of run lengths in Figure 1J (excluding the departure runs). **(C)** Normalized kernel density estimate (KDE) of the wrapped run midpoint in the post-AP period for fly data (re-plotted from the right panel in Figure 1M), random sampling model (N = 300), and Lévy flight model (N = 300).

**Figure S3.**
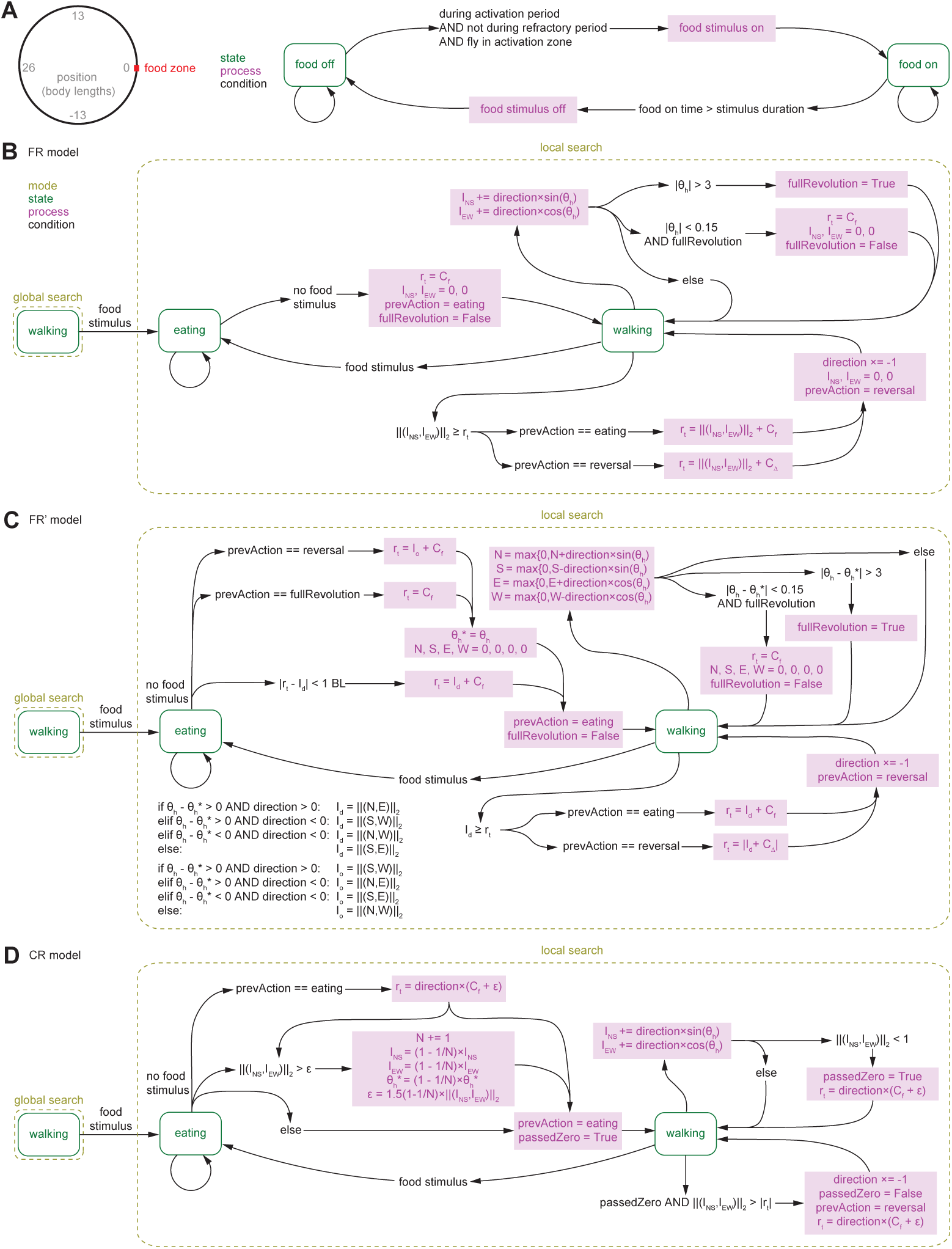
State-transition diagrams describing agent-based odometric integration models of *Drosophila* local search, related to Figure 3. **(A)** Left: Schematic of the simulated environment. The example shown here is for a simulated environment with a single food zone. Right: State transition diagram for the simulated environment. The simulated environment is in either the food on or off state. Transitions between these states, via processes, are determined by the conditions at each timestep of the simulation. See methods for details. **(B-D)** State transition diagrams for the FR model (B), FR’ model (C), and CR model (D). The simulated fly can either be in an eating or walking state, within either a global or local search mode. Transitions between these states and modes, sometimes via processes, are determined by the conditions at each timestep of the simulation. (*rt* = target run length, BL = body lengths, I = Integrator, N = North, S = South, E = East, W = West, θh = heading angle, prevAction = previous action, firstRev = first reversal). The variables Cf, CΔ, CΔ,AP, and CΔ,post-AP, represent a value drawn from the corresponding distribution. See methods for details.

**Figure S4.**
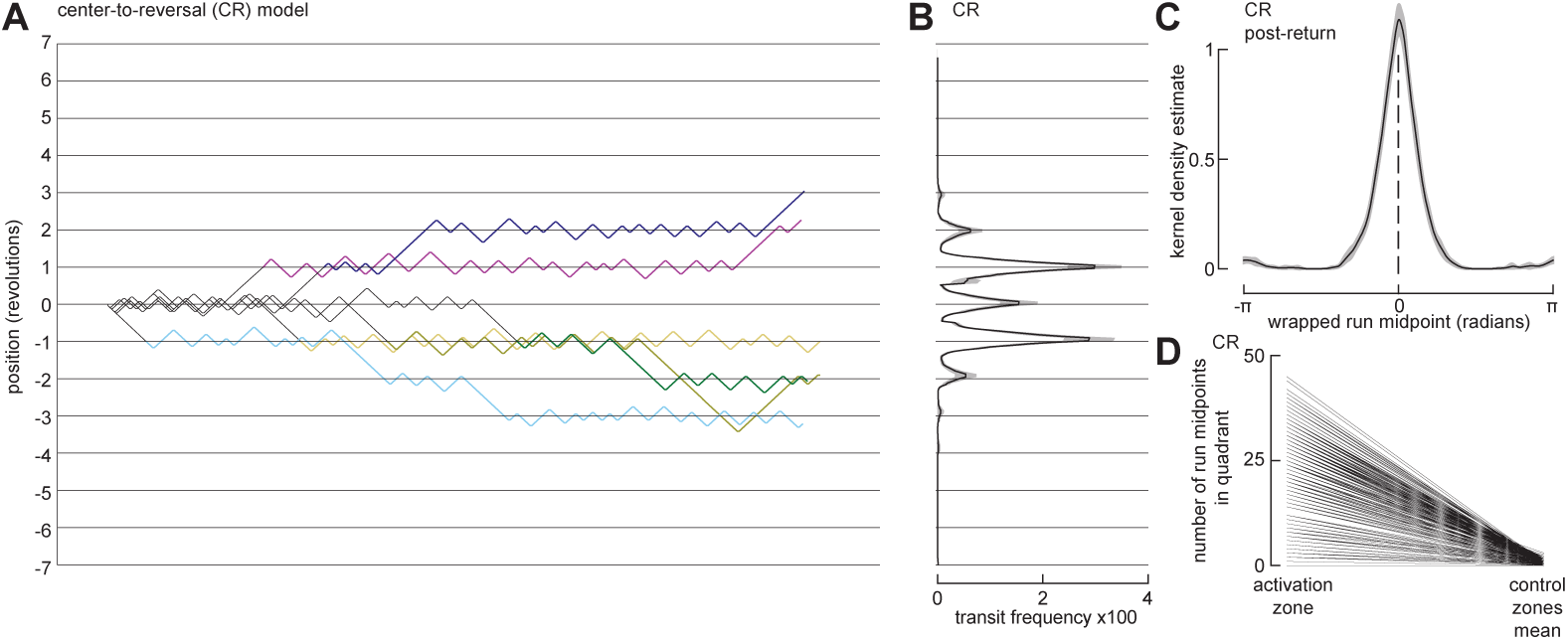
The CR model recapitulates fly re-initiation of local search at a former fictive food site after circling the arena, related to Figure 2. **(A)** As in Figure 2B, for simulations using the CR model. (N = 300). **(B)** As in Figure 2E, for simulations using the CR model. (N = 300). **(C)** As in Figure 2F, for simulations using the CR model. (N = 300). **(D)** As in Figure 2G, for simulations using the CR model. (N = 300).

